# High axial-resolution optical stimulation of neurons *in vivo* via two-photon optogenetics with speckle-free beaded-ring pattern

**DOI:** 10.1101/2022.03.11.484045

**Authors:** Cheng Jin, Chi Liu, Lingjie Kong

**Affiliations:** State Key Laboratory of Precision Measurement Technology and Instruments, Department of Precision Instrument, Tsinghua University,Beijing 100084, China; IDG/McGovern Institute for Brain Research, Tsinghua University, Beijing 100084, China

## Abstract

Two-photon optogenetics has become an indispensable technology in neuroscience, due to its capability in precise and specific manipulation of neural activities. Scanless holographic approach is generally adopted to meet the requirement of stimulating neural emsembles simultaneously. However, the commonly used disk patterns fail in achieving single-neuron resolution, especially in axial dimension, and its inherent speckles decrease the stimulation efficiency. Here, we propose a novel speckle-free, beaded-ring pattern for high axial-resolution optical stimulation of neurons *in vivo*. Using a dye pool and a fluorescent thin-film as samples, we verify that, compared to those with disk patterns, higher axial resolution and better localization ability can be achieved with beaded-ring patterns. Furthermore, we perform two-photon based all-optical physiology with neurons in mouse S1 cortex *in vivo*, and demonstrate that the axial resolution obtained by beaded-ring patterns can be improved by 24% when stimulating multiple neurons, compared with that of disk patterns.

## Introduction

In recent decades, optogenetics has played an increasingly important role in probing functional codes of neurons [1–3]. However, in conventional one-photon stimulation based optogenetics, all neurons in the illumination volume would be excited or inhibited, which makes it complex to decipher the neural network. In order to regulate opsins at single-neuron resolutions specifically, two-photon optogenetics has been demonstrated [4,5].

Considering that opsins distribute on neural membranes, to perform two-photon optogenetics on specific neural ensembles, there are generally two kinds of stimulation strategies: spiral scan approach [6,7] and scanless holographic approach [8,9]. The former approach manipulates neurons by spirally scanning the somas with diffraction limited foci. This method requires lower laser energy, but may introduce latency and jitter. The latter approach generates extended patterns to cover whole somas based on computer-generated holography (CGH). This method can stimulate neurons without temporal delay caused by scanning, resulting in a better temporal resolution. However, for commonly used extended patterns, such as disks, the axial resolution deteriorates linearly with the increase of radii [10]. Thus, it is difficult to maintain single-neuron resolution, especially when multiple extended patterns are required to simulate neural ensembles simultaneously [11].

To improve the axial resolution of extended patterns in scanless holographic approach, temporal focusing (TF) technology is adopted [12]. However, the introduction of grating increases system complexity and loss of laser power. Moreover, the combination of conventional TF and CGH can only improve axial resolution at focal planes. In order to improve axial resolution further in the whole three-dimensional (3D) space, a dual spatial light modulators (SLMs) system is proposed [13], in which the spherical phase is introduced to obtain an extended disk pattern at temporal focus plane. But this method is more complex and produces secondary focus [14].

Besides, as the phase of light field at the target plane is unconstrained in CGH, there are random energy fluctuations on the generated extended patterns, namely speckles, which are more obvious in two-photon optogenetics [15]. To eliminate speckles, various solutions have been proposed. From the perspective of novel CGH algorithms, holographic speckle can be depressed by introducing initial phase at target planes [16], using bandwidth constraint strategy [17,18], adopting zero padding [19], and employing deep learning [20]. But in these methods, parameters should be optimized for specific goals, which increases computational complexity. From the perspective of hardware development, time averaging of multiple irrelevant holograms can also improve the uniformity of holograms [21,22], but this method is not applicable in two-photon optogenetics as it sacrifices temporal resolutions.

Here, we propose a novel scheme to achieve high axial-resolution optical stimulation of neurons *in vivo* via two-photon optogenetics with the beaded-ring pattern. Because the foci in the beaded-ring pattern are separated from each other, no interference leads to speckles, *i.e*. our method is speckle-free. We compare the intensity distribution and axial resolution of different types of stimulation patterns, and demonstrate that higher spatial resolution can be achieved with our novel patterns. Besides, we perform two-photon based all-optical interrogation with neurons in mouse S1 cortex *in vivo*, and demonstrate that the physiological resolution achieved with beaded-ring patterns is better than that with commonly used disk patterns. We expect that the optical stimulation with beaded-ring patterns is a promising strategy for holographic two-photon optogenetics.

## Materials and Methods

### Optical setup

We experimentally characterize two-dimensional (2D) intensity distribution of the generated holographic patterns by the vertical detection method [23] with two-photon excited fluorescence in a dye pool of Sulforhodamine 101 (CAS#60311-02-6, Shanghai Yuanye Bio-Technology). To test 3D intensity distribution when multiple patterns are generated simultaneously, we build a detection system with opposite-facing objectives (Fig. 1). The device used in the excitation path is consistent with that of the vertical detection system [23]. The sample is a custom-made thin film of fluorescent paint (Tamiya Color TS-36 Fluorescent red) on cover glasses. The emission fluorescence is imaged on a CCD (33UX178, DMK) by the objective 2 (25×, 1.05 NA, XLPLN25XWMP2, Olympus), a filter (ET750SP-2p8, CHROMA), and a tube lens (TTL200-A, Thorlabs). Fluorescence intensity characterization of 3D volumes is obtained by moving the objective 1 (25×, 1.05 NA, XLPLN25XWMP2, Olympus) with a piezo-scanning stage, while the objective 2 and the thin film are fixed (the film is at the focal plane of objective 2).

**Fig. 1.**
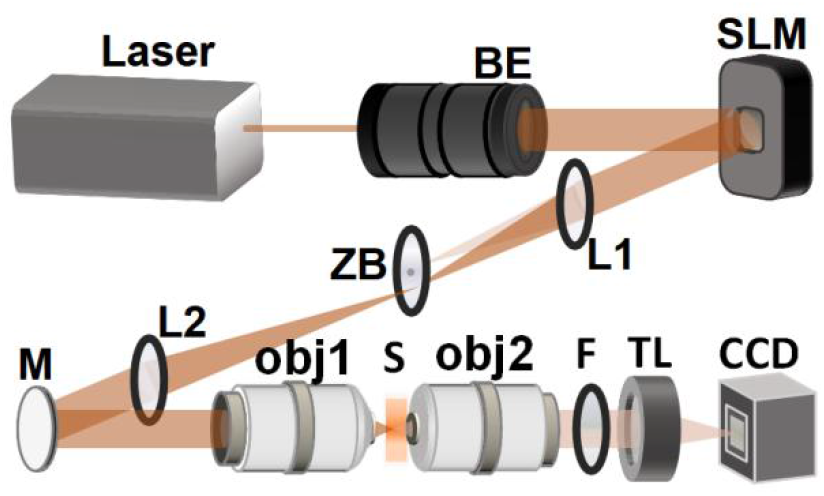
Detection system with opposite-facing objectives. BE: beam expander, ZB: zero-order blocker, L: lens, M: mirror, obj: objective, S: sample, F: filter, TL: tube lens.

To verify the improved performance of our method via two-photon based all-optical physiology *in vivo*, we build a customized two-photon imaging microscope, integrated with a holographic two-photon excitation/inhibition path (similar to the setup in [6]). A SLM (X10468-07, Hamamatsu) is introduced into the excitation path to display holographic extended patterns [8]. The repetition rate of the femtosecond laser for both imaging (*λ* = 920 *nm*) and stimulation (*λ* = 1040 *nm*) is 80 MHz (Chameleon Discovery, Coherent). A hydrogel layer is used for registration between holographic coordinates and two-photon images [6].

### Holographic pattern generation

We use GSw algorithm [24] for holographic pattern generation. Holograms are obtained after 100-times iteration. A disk pattern commonly used for optical simulation covers the whole circle region. In this work, two novel patterns are generated and tested, compared with the disk one. One is an annular pattern, the other is a beaded-ring pattern, which consists of multiple foci evenly spaced along a circle. For annular patterns, the pattern width is 1 *μm*, which is defined as outer radius minus inner radius. For beaded-ring patterns, 8 foci along a circle with a radius of 5 *μm* is used for most experiments, except the patterns in Figs. 2b–2f.

**Fig. 2.**
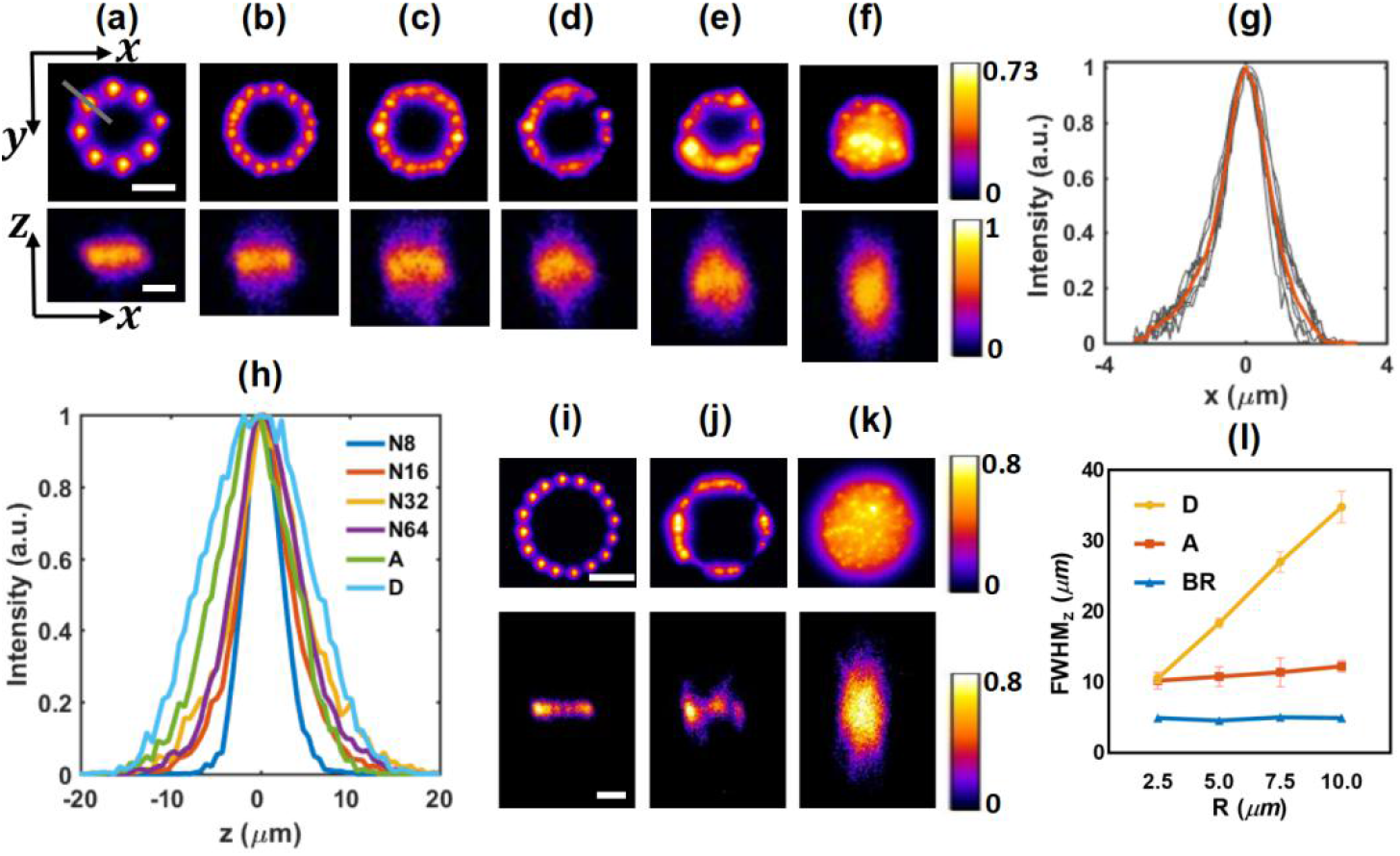
The characterization of generated holographic patterns for a single target stimulation. (a-f) Two-photon intensity distributions in XY (upper) and XZ (lower) section of beaded-ring of 8, 16, 32, 64 foci, annular and disk patterns with 5 *μm* radius, respectively. Scale bar: 5 *μm*. (g) Lateral intensity distribution of each focus in (a) and the average lateral intensity of all 8 foci, labelled with gray and red lines, respectively. (h) Axial intensity distribution of different patterns shown in (a-f). N8: beaded-ring pattern of 8 foci (N16, N32, and N64 have similar definitions), A: annular pattern, D: disk pattern. (i-k) Two-photon intensity distributions in XY (upper) and XZ (lower) section of beaded-ring of 16 foci, annular, and disk patterns with 10 *μm* radius, respectively. Scale bar: 10 *μm*. (l) Axial resolution of the beaded-ring, annular, and disk patterns with different radii. D: disk pattern. A: annular pattern. BR: beaded-ring pattern. Five patterns of different radii for each type are generated, respectively, and the center of each pattern is 20 *μm* away from the origin. The data shown in Fig. 2(l) are the mean and standard deviation of axial resolutions of five patterns.

**Fig. 3.**
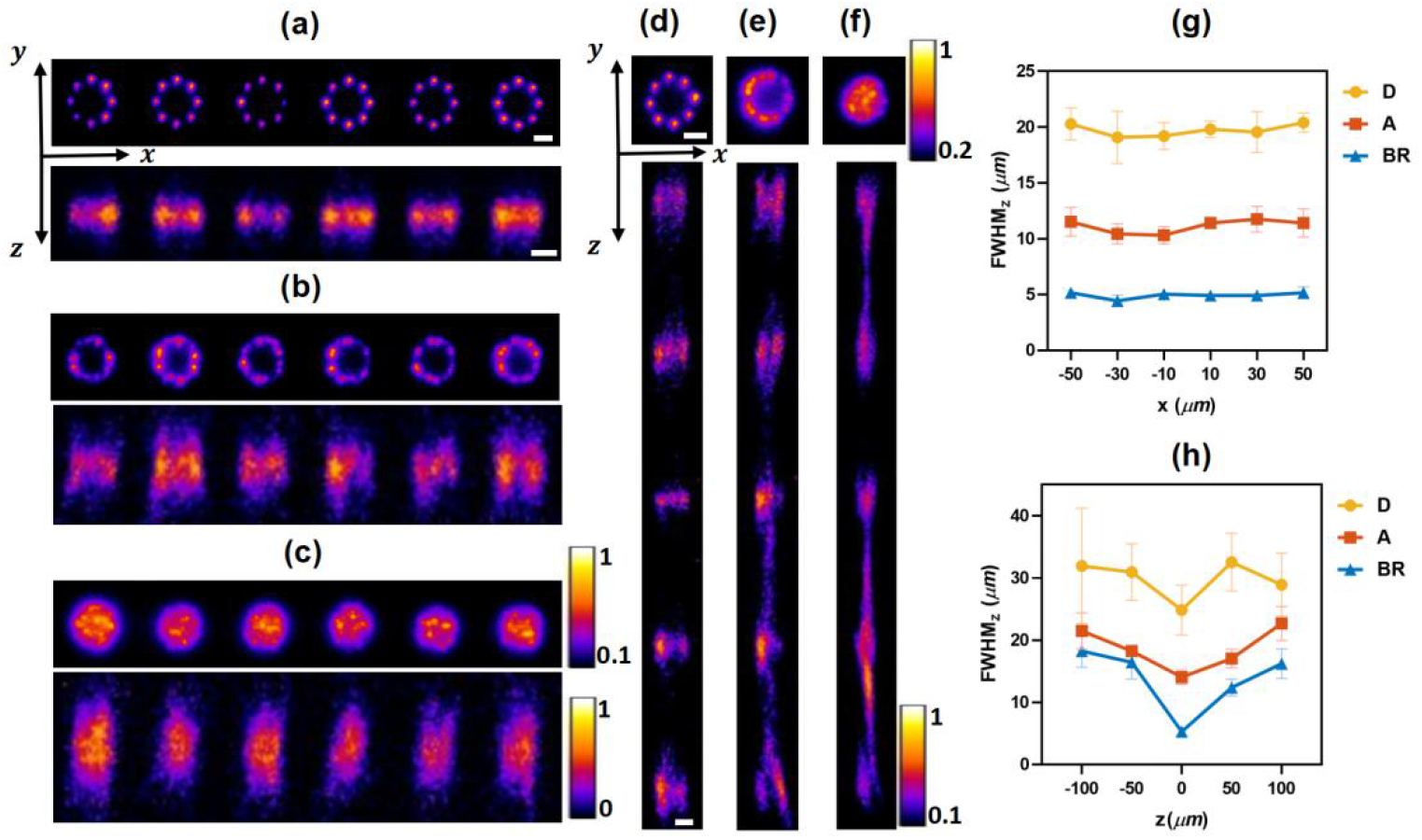
The characterization of generated holographic patterns for multiple targets stimulation in a plane. (a-f) Two-photon intensity distributions in XY (upper) and XZ (lower) section of beaded-ring, annular, and disk patterns, respectively. (a-c) Targets are in the same lateral section (XY), whose centers are from −50 to 50 *μm* with a lateral interval of 20 *μm*. (d-f) Targets are in the same axial section (XZ), whose centers are from −100 to 100 *μm* with an axial interval of 50 *μm*. (g,h) Axial resolution of every pattern in (a-c) and (d-f), respectively. Scale bar: 5 *μm*. All the pattern distribution in Figs. 3(a-f) are obtained under the same excitation power level. D: disk pattern. A: annular pattern. BR: beaded-ring pattern. Five groups of patterns in different distributions (XY or XZ) of each type are generated, respectively, and the distances from patterns of different groups to the center of the field of view are equal for the same distribution. The data shown in Figs. 3(g,h) are the mean and standard deviation of axial resolutions of five group patterns.

In all-optical physiology *in vivo* with two-photon optogenetics experiments, the center of a neuron to be stimulated, which is considered as the center of a target pattern, is manually selected based on the two-photon image at the imaging focal plane. The center positions of patterns at different depths are calculated based on system registration parameters. A series of disk and beaded-ring patterns are generated according to these center positions.

### Sample preparation

All procedures involving mice are approved by the Animal Care and Use Committees of Tsinghua University. We label the neurons in mouse L2/3 of S1 cortex via virus infection with a single integrated ChRmine/GCaMP6m virus [25]. The mice, with chronic optical windows installed after craniotomy, are anesthetized during tests. All-optical physiology tests are performed on mice after 3 weeks of virus expression.

### Data processing

In fluorescence intensity characterization experiments, we integrate intensity within the target pattern at each z-depth to obtain axial intensity distribution curve. The full width at half maximum (FWHM) of the axial intensity distribution curve is defined as the axial resolution of a pattern, no matter for a 2D or a 3D one. The peak value of an axial intensity curve is defined as the maximum intensity, which is used to characterize the energy of a pattern.

To process data in all-optical physiology experiments, a customized Matlab code is used. After summing two-photon signals in a target neuron region and removing background near the neuron, the calcium signal response *F* in each frame is obtained. The change of calcium signal *δF* arised from two-photon optogenetics is defined as the maximum calcium signal caused by stimulation minus the calcium signal just one frame before stimulation. The neural activity response curve is obtained by Gaussian fitting of *δF* at different depths and its FWHM is defined as the physiological resolution of this neuron [6,14].

## Results

### 2D performance characterization of holographic patterns

We firstly use a vertical detection system for the characterization of a single holographic pattern. For a beaded-ring pattern, when 8 foci equally spacing along the circle with a radius of 5 *μm* are generated, the average lateral resolution of these foci is 1.5 *μm* [Figs. 2(a,g)], while the axial resolution is 4.56±0.33 *μm* (mean ±standard deviation). For patterns with the same radius, the axial resolution of an annular pattern is worse (10.8±1.41 *μm*) and that of a disk pattern is the worst (18.36±0.68 *μm*). As shown in Figs. 2(a-f, h), with the increase of focal numbers, the axial resolution gradually deteriorates until it is similar to that of an annular pattern. For disk patterns of different radii, the axial resolution increases linearly with the increase of radius. In contrast, the axial resolution has no significant change with the change of radius for both annular and beaded-ring patterns, and the axial resolution of beaded-ring patterns is always better [Fig.2 (l)].

Next, we characterize multiple holographic patterns with the vertical detection system. The intensity distribution of multiple patterns generated on the same lateral and axial plane by different schemes are shown in Figs. 2(a-c) and Figs. 2(d-f), respectively. As shown in Figs. 2(g,h), higher axial resolution can be achieved with beaded-ring patterns, both for lateral and axial multiple patterns generation. Especially when multiple targets are generated on different axial planes, part of energy is distributed in non-targeted areas for disk patterns [Fig. 2(f)] and annular patterns [Fig. 2(e)]. Significantly, this phenomenon can be avoided in the case of beaded-ring patterns [Fig. 2(d)].

### 3D performance characterization of holographic patterns

We further characterize multiple holographic patterns in 3D [Figs. 4(a-c)], using a detection system with opposite-facing objectives shown in Fig. 1. GSw algorithm is used to generate ten targets of beaded-ring, annular, and disk patterns, respectively. They all have a radius of 5 *μm* and distribute in a 3D space of 100×100×150 *μm^3^*. For beaded-ring patterns, there is almost no energy distribution in non-target areas [Fig. 4(a)]. However, for disk patterns, the energy distribution in non-target areas is high [Fig. 4(c)], which may lead to off-target stimulation in two-photon optogenetics. Besides, the axial resolution of beaded-ring patterns is more than twice better, compared to that of annular and disk patterns [Fig. 4(d)]. In addition, under the same excitation power level, the average signal intensity and signal uniformity of generated beaded-ring patterns are the highest among these three types of patterns [Fig. 4(e)]. All above demonstrate that the beaded-ring pattern has the best capability in spatial confinement and is the most favorable for two-photon optogenetics.

**Fig. 4.**
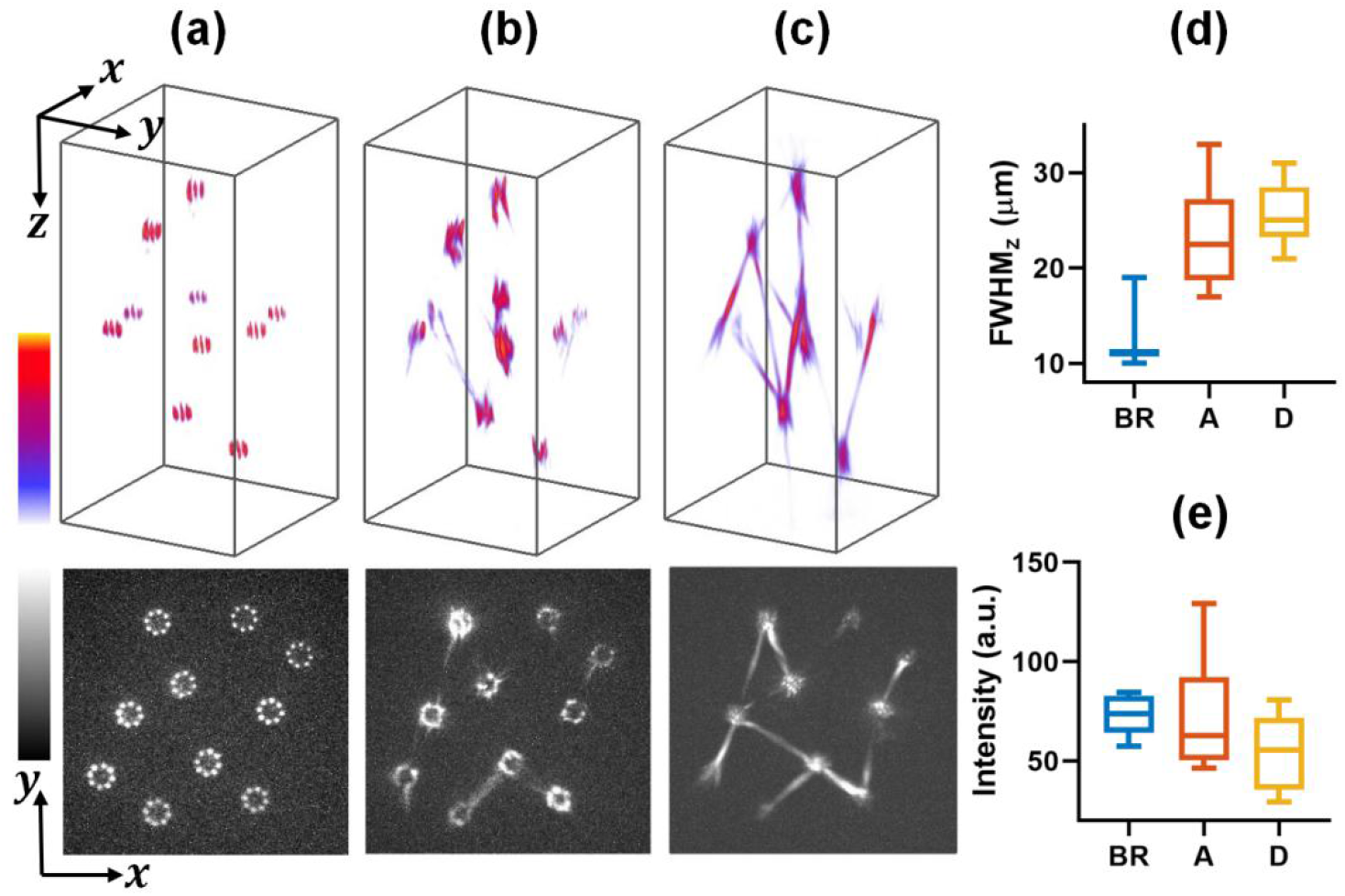
The characterization of generated holographic patterns for multiple targets stimulation in 3D stack. (a-c) Intensity distributions (upper) and maximum intensity projections in XY section (lower) of ten targets with beaded-ring, annular, and disk patterns, respectively. Radius of each pattern: 5 *μm*. Stack size: 120×120×240 *μm*^3^. (d) Statistical analysis of axial resolutions of each pattern in (a-c). (e) Statistical analysis of maximal intensities of each pattern in (a-c). The central mark of each box in (d,e) indicates the median, while the bottom and top edges of the boxes indicate the 25th and 75th percentiles, respectively. BR: beaded-ring pattern. A: annular pattern. D: disk pattern.

### All-optical physiology in vivo with two-photon optogenetics

To verify the performance enhancement of our proposed beaded-ring patterns for two-photon optogenetics *in vivo*, we perform experiments based on all-optical physiology system. Specific holograms are loaded on the SLM to stimulate target neurons with different types of patterns and the corresponding responses of neurons are recorded by two-photon fluorescence imaging of the calcium indicator GCaMP6 [Fig. 5(a)]. By imaging 1 *μm* fluorescent beads with generated holographic patterns in stimulation path, the convolutions of illumination patterns with fluorescent beads are obtained as shown in lower right of Fig. 5(a), which demonstrate the typical intensity distribution of disk and beaded-ring patterns in our all-optical physiology system under the same excitaion power. In the scenario of disk pattern, significant speckles show up, which would definitely lead to low stimulation efficiency. In contrast, a beaded-ring pattern whose foci are evenly distributed on a circle is not interfered by the speckles.

**Fig. 5.**
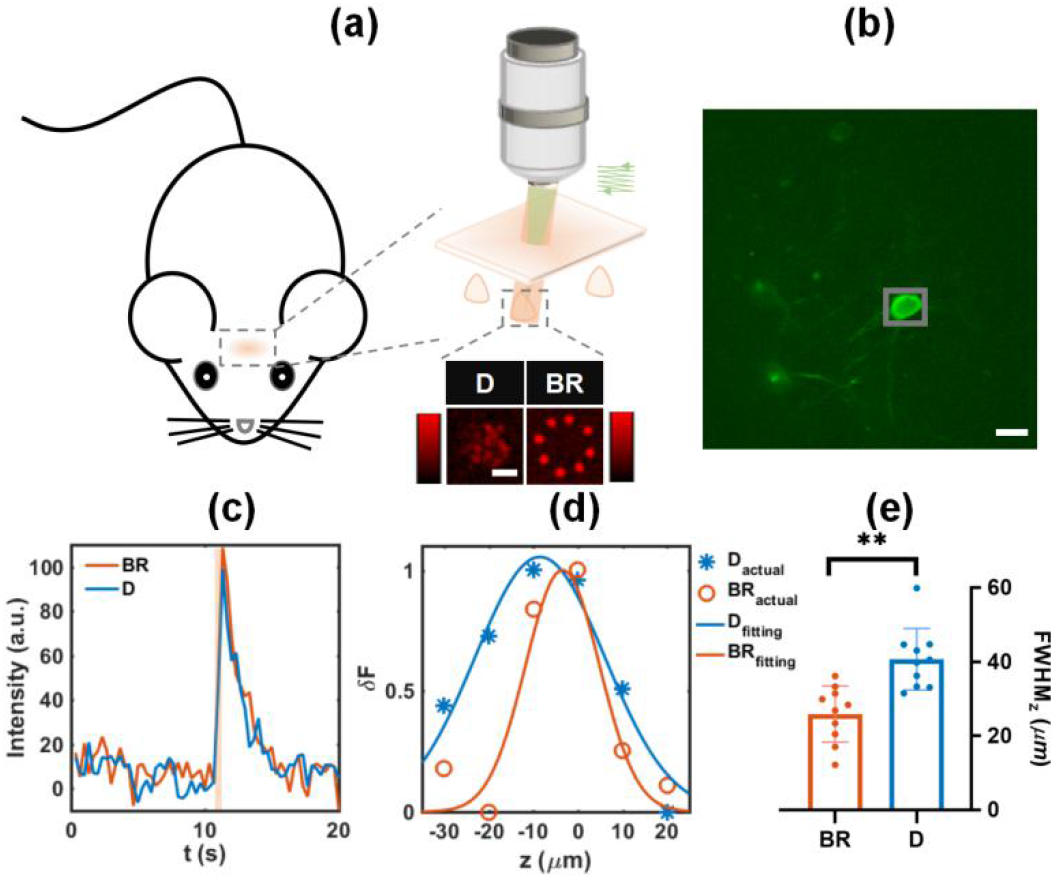
All-optical physiology test with different stimulation patterns on a single neuron *in vivo*. (a) Schematic diagram of the all-optical physiology system. Green: the beam for two-photon calcium imaging. Red: the beam for two-photon optogenetics. Diagram at lower right: the excitation pattern distribution tested by the all-optical physiology system with 1 *μm* fluorescent beads. D: disk pattern. BR: beaded-ring pattern. Scale bar: 5 *μm*. (b) Two-photon image of a neuron in L2/3 of mouse S1 cortex, infected with a ChRmine/GCaMP6m virus. Scale bar: 10 *μm*. (c) Calcium signal of a neuron under different stimulation patterns at the optimal stimulation depth. (d) Changes in calcium signal caused by two-photon optogenetics at different stimulation depths under different stimulation patterns. The fitted response intensity curves are shown with solid lines. (e) Statistical analysis of axial resolutions under different stimulation schemes. Data are from 10 neurons in 3 mice. **p=0.0017, ratio paired t test.

We achieve the location of neurons with two-photon imaging mode. For a target neuron in an imaging field-of-view [Fig. 5(b)], two-photon optogenetics based on disk and beaded-ring patterns are employed to stimulate the neuron respectively, and the photo-stimulation duration is 100 *ms*. In order to restore opsin after each stimulation, we set a dark interval of at least 60 seconds between two tests. To avoid pseudo activation of neurons by the imaging beam, the imaging power is reduced as much as possible so that the calcium baseline of neurons can hardly be seen when only two-photon imaging is performed. The powers of stimulation and imaging vary slightly due to different expression levels of opsins and calcium indicators among different neurons in different mice.

The stimulation depth where the maximum *δF* is obtained among a series of stimulation depths is defined as the optimal stimulation depth. We find that a target neuron can be stimulated by both types of patterns at their optimal stimulation depths with the same excitation power [Fig. 5(c)]. *δF* decreases gradually when deviating from the optimal stimulation depth, and the decrease rate is faster for the case with a beaded-ring pattern than that with a disk one [Fig. 5(d)]. By testing and analyzing physiological resolutions [6,14] of 10 neurons with single neuron stimulations, we find that the axial resolution under optical stimulation with beaded-ring patterns (25.91±7.57 *μm*) is significantly better than disk patterns (40.68±8.33 *μm*), as shown in Fig. 5(e). The improvement is as much as 36.3%, which indicates the capability of optical stimulation by beaded-ring patterns at single-neuron resolutions.

Next, we perform two-photon optogenetics stimulation and two-photon calcium signal imaging experiments on multiple neurons simultaneously. As shown in Fig. 6(a, left), two neurons to be stimulated are selected in a same field-of-view, limited by the available laser power. Then, beaded-ring and disk patterns matching these two neurons are used for stimulation [Fig. 6(a, right)], with duration of 100 *ms*. Two neurons stimulated at the same time are defined as a group. Compared with disk patterns, the average axial resolution of neurons in a group stimulated by beaded-ring patterns is increased by 24.27% [Fig. 6(b)], from (38.24±6.55 *μm*) to (28.96±8.19 *μm*). The mean peak responses of beaded-ring patterns are not significantly different from that of disk patterns [Fig. 6(c)], which shows that the beaded-ring pattern has comparable excitation efficiency with the disk one at the optimal stimulation depth.

**Fig. 6.**
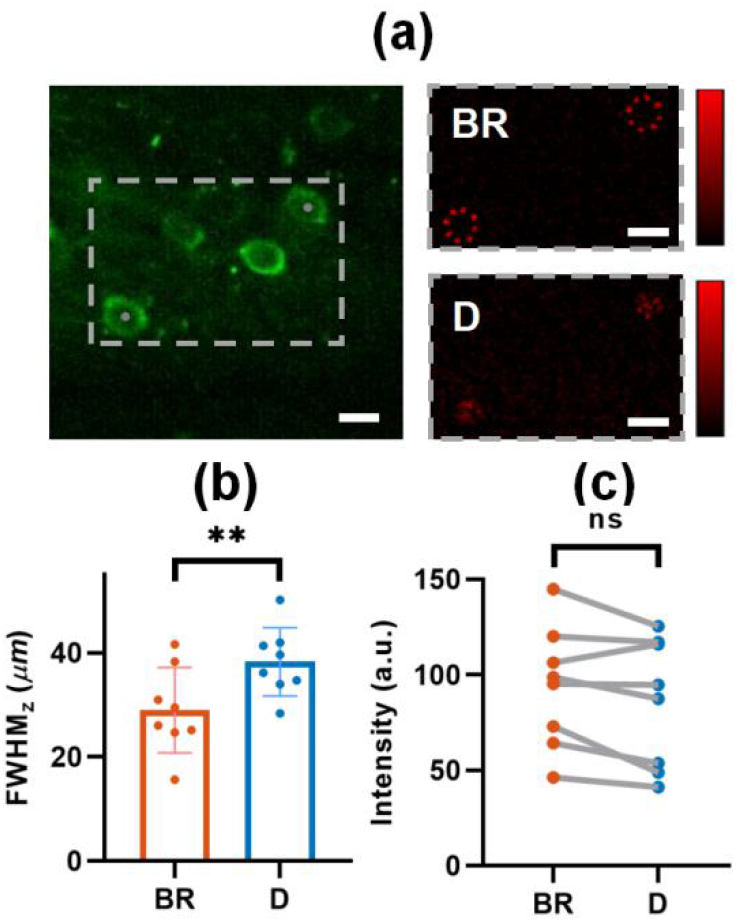
All-optical physiology test with different stimulation patterns on multiple neurons simultaneously *in vivo*. (a) Left: a typical two-photon image of neurons in L2/3 of mouse S1 cortex, infected with a ChRmine/GCaMP6m virus. Locations of gray spots are the center of two excitation patterns. Right: the distribution of excitation patterns to stimulate target neurons. D: disk pattern. BR: beaded-ring pattern. Scale bar: 10 *μm*. (b) Mean axial resolutions of calcium signals of multiple neurons under different stimulation schemes. Data are from 8 groups and each group has two neurons to be stimulated. **p=0.0086, ratio paired t test. (c) Mean peak responses of multiple neurons at their optimal stimulation depths under different stimulation schemes. Data are from 8 groups and each group has two neurons to be stimulated. ns: not significant, p=0.0638, ratio paired t test.

## Discussion and Conclusion

In this paper, we propose a novel speckle-free, beaded-ring pattern for holographic two-photon optogenetics *in vivo*. Compared with conventional disk patterns, higher axial resolution can be achieved with beaded-ring patterns. Meanwhile, the axial resolution of a beaded-ring pattern is not affected by the increase of radius. When multiple patterns are generated simultaneously in 3D space, the laser power of beaded-ring patterns can be restricted in target regions, while laser power often flows to non-target regions when disk patterns are generated. Through all-optical physiology test *in vivo*, we demonstrate that the physiological resolution in axial dimension can be improved by as much as 24.27% by using beaded-ring patterns compared to that with disk patterns. We expect a broad application of our proposed beaded-ring patterns for holographic two-photon optogenetics.

## Acknowledgments

The authors thank Yang Lin and Kuikui Fan for helps on bio-sample preparation.

## Data availability

All data presented in this paper are available upon reasonable request from the corresponding author.

